# Steep difference between ecotypes and intermediate phenotype depression for shell trait integration in a marine snail hybrid zone

**DOI:** 10.1101/054569

**Authors:** Carlos Garcia

## Abstract

Multivariate analyses of phenotypic integration for a set of characters provide information about biological systems that cannot be obtained in univariate studies of these characters. We studied phenotypic integration for seven shell measures across the phenotypic gradient in a hybrid zone of the marine snail *Littorina saxatilis* in Galicia, NW Iberia. We found clear differences in the degree of integration between the two ecotypes involved in the hybrid zone, likely related to differences in the strength of natural selection acting on the snails' shells in each ecotype's habitat. We found also evidence of a decrease in integration in the phenotypically intermediate, hybrid snails, consistent with hybridization resulting in a release of multivariate variation and increased evolvability. Across the phenotypic gradient, decreases in overall integration tended to be accompanied by increases in some measures of modularity, but the latter did nor reflect high correlation structure. The increases occurred only in a proportional sense, correlations among modules tending to decrease faster than within modules for low overall integration tiers. Integration analyses based on non partial and partial correlations tended to produce contrasting results, which suggested hierarchical sources of shell integration. Given that the two ecotypes could have differentiated in situ according to a parapatric model, our results would show that changes in integration can occur in a short evolutionary time and be maintained in the presence of gene flow, and also that this gene flow could result in the hybrid release of multi character variation.

## INTRODUCTION

Because natural selection acts on whole phenotypes (Sober 1984), the fitness of organisms depends on combinations of different characters, and studies considering individual traits separately may be incomplete or even misleading (Pigliucci 2003, Blows 2007). New properties emerge when sets of different characters are considered jointly, one of the most relevant being integration, the cohesion among traits that results from the interactions between the different biological processes producing the phenotype (Pigliucci 2003; Klingenberg 2008, Murren 2012). Integration may be studied from the point of view of the developmental, regulating or functional processes resulting in the observed phenotypes (Klingenberg et al. 2014, Armbruster et al. 2014, Meyer et al. 2014), but most often, integration studies focus on the observed phenotypic correlations among traits (Pigliucci 2003, Monteiro and Nogueira 2010; however, these correlations may not accurately reflect the physiological or developmental sources of the relationships between characters, so that the more restrictive term “statistical integration” has been proposed for such studies; see Armbruster et al. 2014), the phenotypic correlation matrix playing a central role in the study of the evolution of complex traits. Since correlations between characters may constrain the evolution of populations in the future, measures of integration and its stability in time may be key in evaluating the evolvability of populations (Schlichting 1989, Wagner and Altenberg 1996).

At the same time, integration may itself be seen as a character under genetic control, whose value is the consequence of the action of natural selection in the past (Olson and Miller 1958, Van Valen 1965, Cheverud 1996a). The study of present integration patterns could thus provide valuable information about the evolutionary history of characters and populations. However, the relationship between evolutionary past and present integration is not completely understood (Gerber 2013, Watson et al. 2014, Leslie et al. 2015, Goswami et al. 2015), and generalizations are difficult, given the relatively few results available about this problem (Klingenberg et al. 2014). It is in principle expected that correlational selection for two characters tends to generate a genetic correlation of the same sign as the selection gradient (Lande 1980, Cheverud 1996b, Sinervo and Svensson 2002, McGlothlin et al. 2005, Roff and Fairbairn 2012, but see Armbruster et al. 2014). Thus, sets of characters having experienced strong correlational selection in the past (as could be the case if they were functionally related) would be highly integrated, and specialist species, adapted to live in a narrow range of environments, would be more integrated for the relevant character sets than generalists (Berg 1960, Ordano et al. 2008, Eroukhmanoff and Svensson 2008, Hermant et al. 2013, Gómez et al. 2014).

Measuring integration is specially interesting in the context of hybridization processes. Comparing integration levels may give clues about both the genetic differences between the hybridizing populations and the evolutionary potential of the resulting hybrids. Integration levels in hybrids could decrease (as observed by Parsons et al. 2011, Saetre 2013, Selz et al. 2014) due for example to the disruption of epistatic interactions or gene regulation networks (Landry et al. 2007, Tulchinsky et al. 2014). The loosening of trade-offs and constraints present in the parent populations would boost genetic variance and permit hybrids to evolve to reach unoccupied adaptive peaks (Klingenberg et al. 2004, Gross and Rieseberg 2005, Mallet 2007, Campbell et al. 2009, Eroukhmanoff et al. 2013, Nichols et al. 2015). But, in principle, changes in integration could occur in any direction: hybrids could show higher integrations (as observed by Murren et al. 2002) because genes from different parents could remain linked (Wagner et al. 2007) or because of heterosis for developmental stability (Renaud et al. 2012).

Because the relationship between population evolutionary history and integration seems to be complex, it is important to increase the information about how real world evolutionary mechanisms and contexts influence phenotypic integration. This information would be more valuable the better understood these mechanisms and contexts are. Here we analyze the changes in integration along the shell phenotype gradient in a hybrid zone of the seashore snail *Littorina saxatilis* that has been the object of evolutionary studies for more than twenty years.

### The Galician Littorina saxatilis hybrid zone

The most wave-exposed shores of Galicia, NW Iberia, are inhabited by two ecotypes of the marine snail Littorina saxitilis (see Rolán-Alvarez, 2007 and Johannesson *et al.*, 2010 for general reviews). One occurs on the drier, upper shore and has large, thick, ridged and banded shells (RB, “crab” ecotype) with narrow apertures, which result in resistance to both predation by the crab *Pachygrapsus marmoratus* (Fabricius, 1787) and desiccation. The other is in the lower and wave-exposed shore, and has small, thin, smooth and unbanded (SU, “wave” ecotype) shells, with wide apertures and low spired forms, which give them resistance to wave action. Gene flow between ecotypes occurs (Galindo et al. 2013) through a hybrid zone on the mid shore. Divergent selection (Rolán-Alvarez *et al.*, 1997, Cruz and Garcia 2001, Cruz *et al.*, 2004a), assortative mating (Conde-Padin *et al.*, 2008) and habitat preference-guided movements (Cruz *et al.*, 2004b) operate against this gene flow to maintain the hybrid zone stable. These populations have been proposed as a case of parapatric ecological speciation (Quesada *et al.*, 2007, Butlin et al. 2014), which makes them a particularly interesting object of study in the present context. The speciation transition is the most important level at which to examine any changes in phenotypic integration and covariance structure, because it is in this transition that the effects of microevolutionary processes can shape macroevolutionary patterns (Steppan, 1997).

### Measuring integration

The most widespread measure of overall integration is the variance of eigenvalues of the correlation matrix (Wagner 1984; Pavlicev et al 2009; see Armbruster et al. 2014 for a review of integration indexes): in the case of maximum integration, all measures vary jointly, so that a single eigenvector explains all the variation (all eigenvalues but one have zero value and there is a large variance between eigenvalues); in the case of minimum integration, all (standardized) variables are independent and make equal contributions to the total variation, all eigenvalues thus being equal and greater than zero and the variance between eigenvalues being zero. Integration may occur and be measured at many phenotypic scales. Measures made at the whole organism level could be difficult to interpret and make limited sense, as different character sets in the same organism may be integrated to different extents (Pigliucci 2003, May 2006, Ordano et al. 2008). Analyses focusing on particular biological functions or character sets are expected to be more useful (Pigliucci 2003). However, even measures of the overall integration in a given character set may be misleading if the set is composed of different modules, i. e., it may be decomposed into subsets of characters having tight relationships with other characters in the same subset but looser ones with those in different subsets (Cheverud 1982, Wagner 1996, Murren 2012). In fact, this modular organization is pervasive at all levels of biological organization, from the genetic to the developmental, anatomical and behavioral (Müller 2007). Because eigenvalue variance would be affected by module size and between-module correlation (Pavlicev et al. 2009), a character set with weak correlations between modules could show limited overall integration, despite having a well defined and biologically relevant modular correlation structure (Conner et al. 2014; see Figure 1). Thus, exploratory integration analyses should measure the degree of modularity in addition to overall integration (Klingenberg 2009, Diggle 2014). The Newman and Girvan's (2004) coefficient is a commonly used measure of the degree of modularity. It is based on comparing the density of correlations (or, in a more general network context, “links”) among variables (more generally “nodes”) in the same module relative to that between variables in different modules:

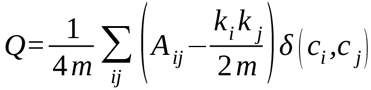

where *A_ij_* = 1 if a link connects variables *i* and *j* and *A_ij_* = 0 otherwise, *k_i_* is the degree of (number of links connected to) variable *i, m* is the total number of links in the network, and the function 5 yields one when variables *i* and *j* are in the same module and zero otherwise.

**Figure 1.**
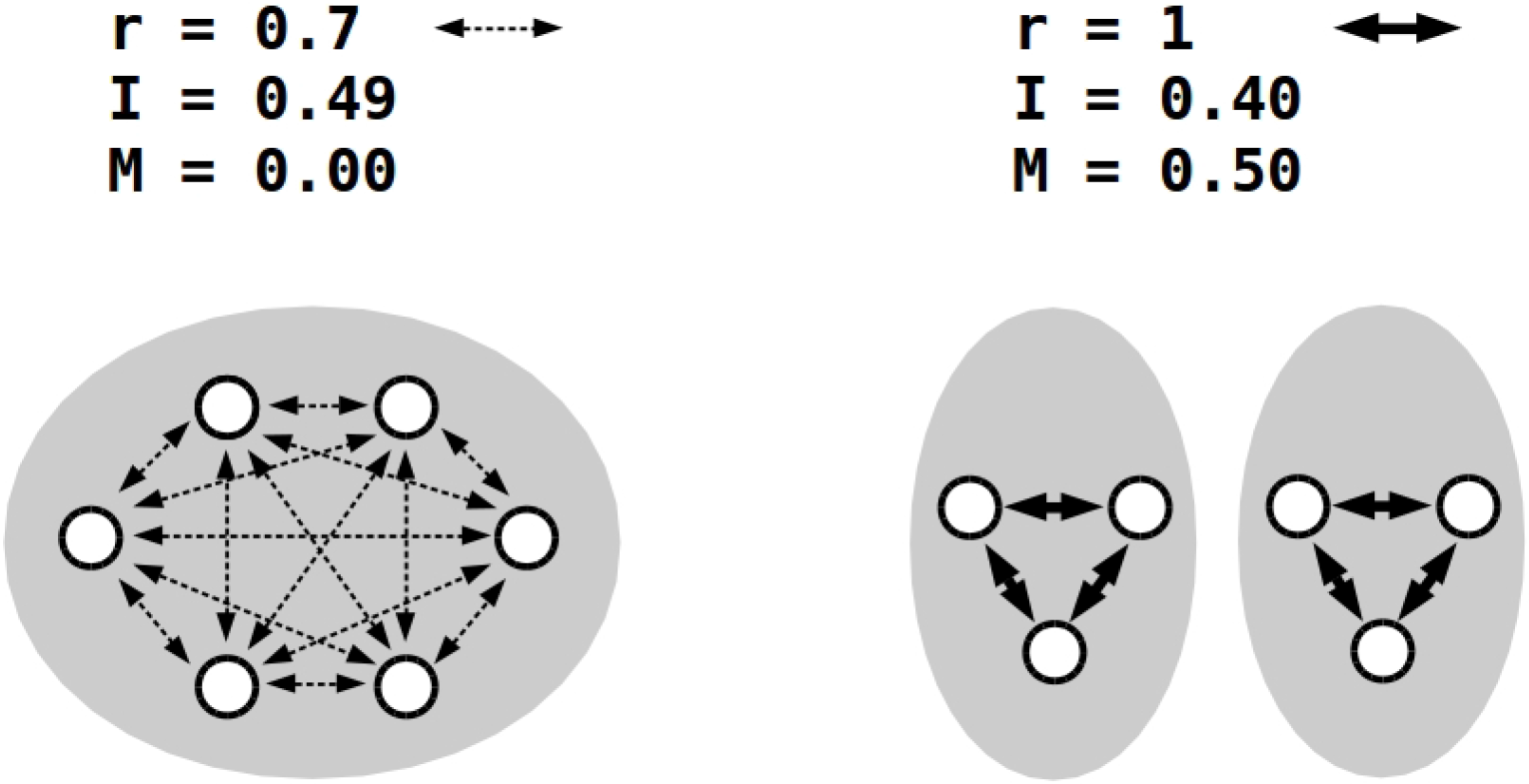
Overall integration versus modularity. Left, six variables (white circles) in a dense web of pairwise correlations are grouped in a single cluster (grey ellipse); they show high overall integration without modularity. Right, six variables have strong correlations within two separated clusters. They make a clear-cut network structure and show high modularity but less overall integration than in the left side case. r, pairwise correlation between variables; I, overall integration as measured by the eigenvalues variance (all variables had the same variance); M, Newman and Girvan modularity coefficient.

Modularity, besides overall integration, might have an impact by itself on the evolutionary potential of populations, as it has been speculated that populations having their characters grouped in clearly defined modules could adapt in a short time to environmental change by producing new combinations of these whole modules, which could result in large scale, integrated phenotypic changes (Wagner and Altenberg 1996, Mezey et al. 2000, Muller 2007, Pavlicev and Hansen 2011). Such nearly independent character modules have been found in morphology studies by Bissel and Diggle (2010), Mallarino et al. (2011), Larouche et al. (2015) and Powder et al. (2015).

## Material and Methods

### Snail Samples and phenotypic measures

The shell data in the present work had been recorded in a between-shore levels reciprocal transplant experiment to measure natural selection on snail shell shape (Cruz et al. 2004a). The populations studied were in Portocelo (41° 59’ N, 8° 53.’ W) and Corrubedo (42° 32’ N, 9° 2’ W), within 63 Km of each other. In each locality, samples of 125 adult snails were taken at five sampling points in a linear transect perpendicular to the shore line: in the lower shore, in the lower shore-mid-shore border (where the frequency of upper shore phenotypes became approximately less than 5%), in the mid-shore (where the frequency of upper and lower shore phenotypes was similar), in the midshore-upper shore border (where the frequency of typical lower shore phenotypes became approximately less than 5%), and in the upper shore. The transects covered most of the of the shore band occupied by the lower shore ecotype, but only a section of the wider upper ecotype's band.

These two data samples were appropriate for the present analysis because they covered a wide range of phenotypes and shore levels, and included significant proportions of intermediate phenotypes. Seven linear measurements (Fig. 2 top) were made on every shell. These variables were combined in a discriminant function ( with coefficients -17.361, 5.268, 1.030, -14.766, -1.096, 3.967 and 19.194 for measures 1 to 7) separating snails from the two extreme shore levels, and the individuals' ranks for the function value were used to divide the sample in each locality into 9 discriminant function tiers of the same size (613/9 and 609/9, i. e., 68 and 67, with rests of 1 and 6 snails added to the ninth tier in Portocelo and Corrubedo; note that not all 625 snails initially sampled in each location produced usable shell measures). The pure ecotype distributions had limited overlap, and the phenotypic distribution of hybrids did no exceed that of these pure ecotypes (Figure 2, middle). Thus, as expected in systems driven by divergent selection (Stelkens et al. 2009) no evidence for transgressive segregation was found in this hybrid zone. Shell morphology was narrowly related to both ecotype category and shore level (Figure 2, bottom). Using snail linear measurements made it possible to take advantage of these already recorded data sets. In any case, very similar results may be obtained in integration and modularity studies using linear measurements or landmark-based geometric morphometrics (Jojic et al. 2012).

**Figure 2.**
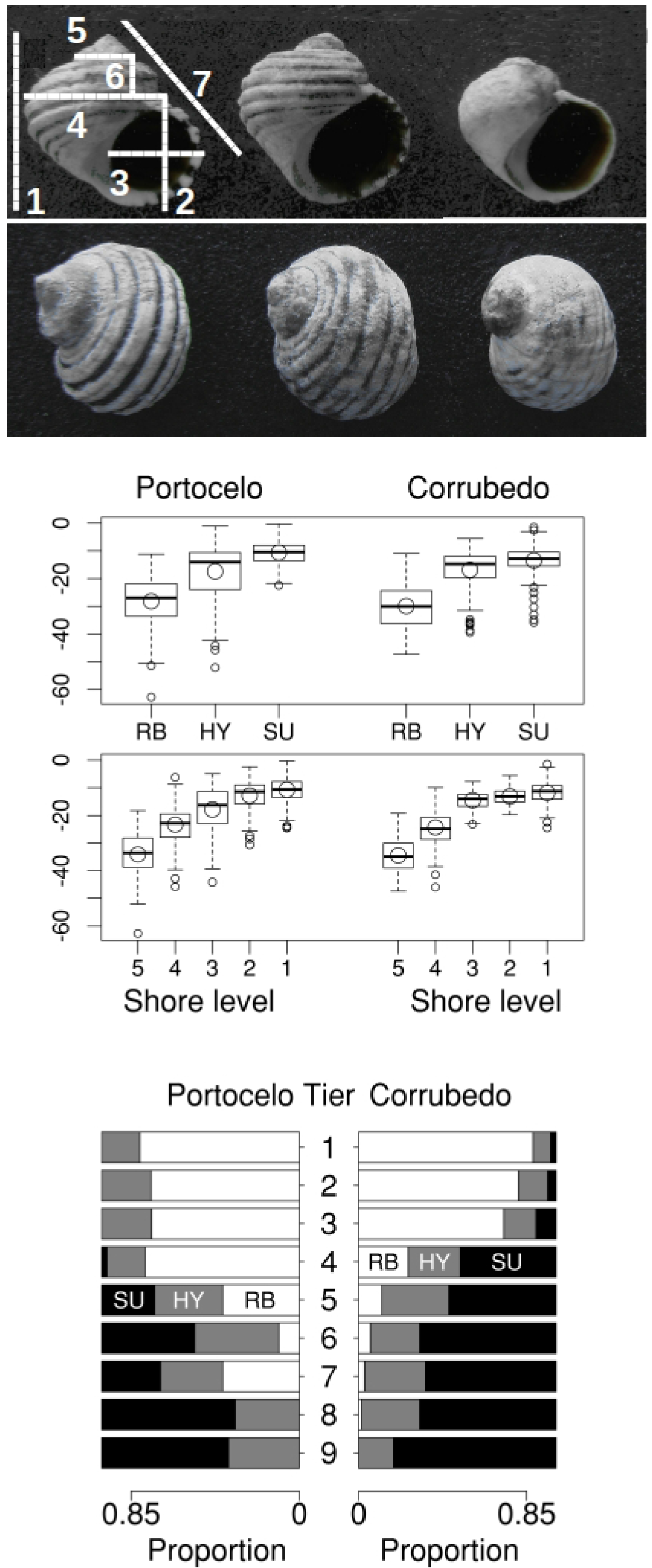
Top, snails from the RB upper-shore ecotype (left), SU lower-shore ecotype (right) and phenotypically intermediate (“hybrid”, HY, center). An smaller than average sized RB and a larger than average sized SU snail are shown to ease shell shape comparisons. The seven shell measures used are indicated on the RB shell. Middle, boxplots of the discriminant function values for the three ecotypes and the five sampled shore levels (5, upper shore; 1, lowest shore) in the two localities. Medians and means are shown as bold horizontal bars and large circles respectively. Bottom, proportion of RB, HY and SU ecotypes in the nine discriminant function-defined phenotypic tiers in the two localities. Individuals were tagged as hybrids when they had bands but not ridges, or the reverse, or had incomplete sets of both features (Johannesson and Tatarenkov 1997).

### Data Analysis

Different measures of integration and modularity were taken in each of the eighteen data sets (9 tiers × 2 localities). The overall integration of the seven shell variables was measured as the variance of their eigenvalues (transformed so that their sum was equal to one in all sets; Wagner G. P. 1984, Pavlicev et al. 2009) and as the average of all absolute correlations (Cane 1993).

A first attempt to analyze shell modularity based on (non-partial) correlations used procedures for community detection in networks implemented in the R package *igraph* (Csardi and Nepusz 2006). However, these procedures, while typically fast and efficient in the analysis of large networks, may face a “resolution limit” (this is specially the case for procedures based on modularity, see Fortunato and Barthélemy 2007) when modules are very small, as is necessarily the case for the seven variables problem studied here. In fact, no communities were detected, so that the alternative procedure BoCluSt, showing a good performance for small to moderate sized data sets (Garcia 2014a) was used in its place. BoClust is a stability-under-resampling procedure that carries k-means clustering analyses of resampled data sets and determines which of the successive cluster-module partitions results in more stable variable allocations, i. e., which is the set of variables-to-modules allocations showing the least variance across resamples. This variance constitutes a natural base to measure the degree of modularity in a set of variables, as highly modular data sets would consist in clearly defined modules-communities resulting in very stable variable allocations and low variances. We calculated the modularity in each snail phenotypic tier as one minus the variance of the variable's allocations in the best module partition in that tier. BoCluSt was very effective in the identification of stable module partitions, to the point of identifying more than one partition with the maximum stability (i. e., a zero allocation variance, and thus an equal to one modularity) in several tiers. To identify the best partition among these, additional BoCluSt runs were made under the “thinning” option, which generates resamples less than full tier sample size (resample size 15 was used for the resamples). The resolution of the analysis is thus reduced so that it becomes harder to get maximum support partitions and the previous ties are resolved. Additionally, the *igraph* procedure was used to calculate the Newman and Girvan modularity coefficient for the BoCluSt identified module partitions. The Cheverud's index was used as a an alternative measure of modular structure. It is calculated as one minus the geometric mean of the correlation matrix eigenvalues (Cheverud et al. 1983). This would be sensible to the “parcellation” among characters (Wagner 1984), i. e., the division of characters sets into subsets. Finally, as a more intuitive measure of this structure, we calculated the ratio: average correlation within BoCluSt identified clusters / average correlation between clusters.

To study the possible existence of overlapping and hierarchical patterns of integration in these data (Wagner and Altenberg 1996, Hansen et al. 2003), we complemented the previous community analyses by calculating the conditional independence matrices of graph modeling analysis (Magwene 2001). We calculated the partial correlations between variables ρ_ij_ and the strength of the edges between them, i. e., the information in variable X_i_ on X_j_ and vice versa conditional on all remaining variables as

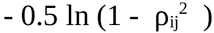

(Whittaker 1990). We discarded all edges too close to zero using the “edge exclusion deviance” statistic (Whittaker 1990):

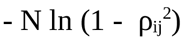

where N is the number of observations. All edges with deviance smaller than 3.84 (the value corresponding to 5% probability in a Chi - square distribution with 1 d. f.) were taken as zero. We obtained additional measures of overall integration based on these analyses. First, the average partial absolute correlations for the seven shell variables, and second, the networks' weighted connectivity, defined as the average strength of edges per node (Proulx et al. 2005).

BoCluSt does not take link weights (such as edge strengths in this case) as input, but the Louvain procedure (Blondel et al. 2008) in *igraph* was able to detect communities when using edge strengths, so that these communities served for the calculation of edge strength-based Newman and Girvan modularities. The better results for the Louvain procedure were likely due to the use of the edge deviance statistic resulting in “sparser” networks with fewer edges, which usually eases community detection (Fortunato 2010). The edge strenghts were used also to calculate the ratio of average edge strengths within / between Louvain identified clusters. No corrections for number of variables or observations were made for any of the above statistics because all comparisons involved the same shell variables and almost equal data sizes.

### Natural selection analysis

We compared the phenotypic correlations and estimates of natural correlation selection obtained in the analysis of the snail data used in the present work. These data had been subjected to multiple logistic regression analysis to estimate direct and divergent selection pressures in the hybrid zone (Cruz et al. 2004a). This regression had considered measures 2 to 7 as relative to measure 1. To make possible a direct comparison between natural selection and phenotypic integration results, in the present work regression model we replaced the relative shell measures with their non-relative equivalents. The measures were log-transformed, standardized within locality and pooled into a single data set (1107 observations; this number is less than that of measured snails because not all these snails provided recapture data useful for the reciprocal transplant experiment). The regression model (*glm* function, R Core Team 2015) included the transplant destination, the shell measures, the destination x measures crossproducts, to estimate divergent selection acting on each measure, the measure *i* x measure *j* crossproducts, to estimate correlational selection, and the destination x measure *i* x measure *j* triple crossproducts to estimate divergent correlational selection (56 independent variables in total). This was complemented with a stepwise regression of the same model (*glm* and *sep* functions, direction = “forward”; R Core Team 2015). To evaluate the degree of model adjustment to the data in this logistic regression, we used the McFadden's pseudo-R squared (McFadden, 1974; Hosmer et al. 2013), defined as

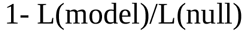

where L(model) is the log likelihood value for the fitted model and L(null) is the log likelihood for a null model including only an intercept as predictor. In contrast with the Akaike information criterion (AIC), this measure does not include a penalization proportional to the number of independent variables fitted. Such penalization would have been irrelevant for the comparisons made here, involving models with equal numbers of variables. Instead, the McFadden's pseudo-R squared measures the likelihood of the fitted model relative to a null, intercept-only model. This relative measure is more appropriate for comparing the analyses made here, which used different data sets (phenotypic tiers).

## Results

### Cluster analysis

The pattern of clustering was different in the two extremes of the phenotypic range (Figure 3). In the upper-shore-like tiers, and with only one tier as an exception, the clustering resulted in the serial separation of variables number 6, 5, 2, 3, and 4 in individual clusters, whereas the order in the lower shore-like tiers was, again with a single exception, 5, 6, 3, 2 and 4. The exceptions in the two extremes were of a rather minor significance: in both, variables 2 and 3, instead of separating sequentially in the third and fourth clustering cycles, formed together a new cluster in the third cycle and separated from each other in the fourth. The differences in clustering pattern show that the two ecotypes have somewhat different correlation structures for these shell shape characters. In the intermediate tiers, the patterns were not simply midway between those in the extreme tiers. For example, in both localities some tiers had variable 4 separated from the main cluster already in the third clustering round, in contrast with the extreme tiers in which this variable always separated last. In addition, the intermediate tiers had more heterogeneous patterns, which suggests a less well-defined correlation structure. Many tiers had more than one clustering with maximum BoCluSt support. The modular structure for these seven measures was therefore hierarchical: the identified clusters were composed of clearly defined sub clusters. In what follows, the BoCluSt, Newman and Girvan measured modularities and within and between clusters average correlations are based on the more interesting and informative three-cluster partitions (as seen in Figure 3, all six-cluster partitions resulted in five single-variable clusters and a cluster including variables one and seven).

**Figure 3.**
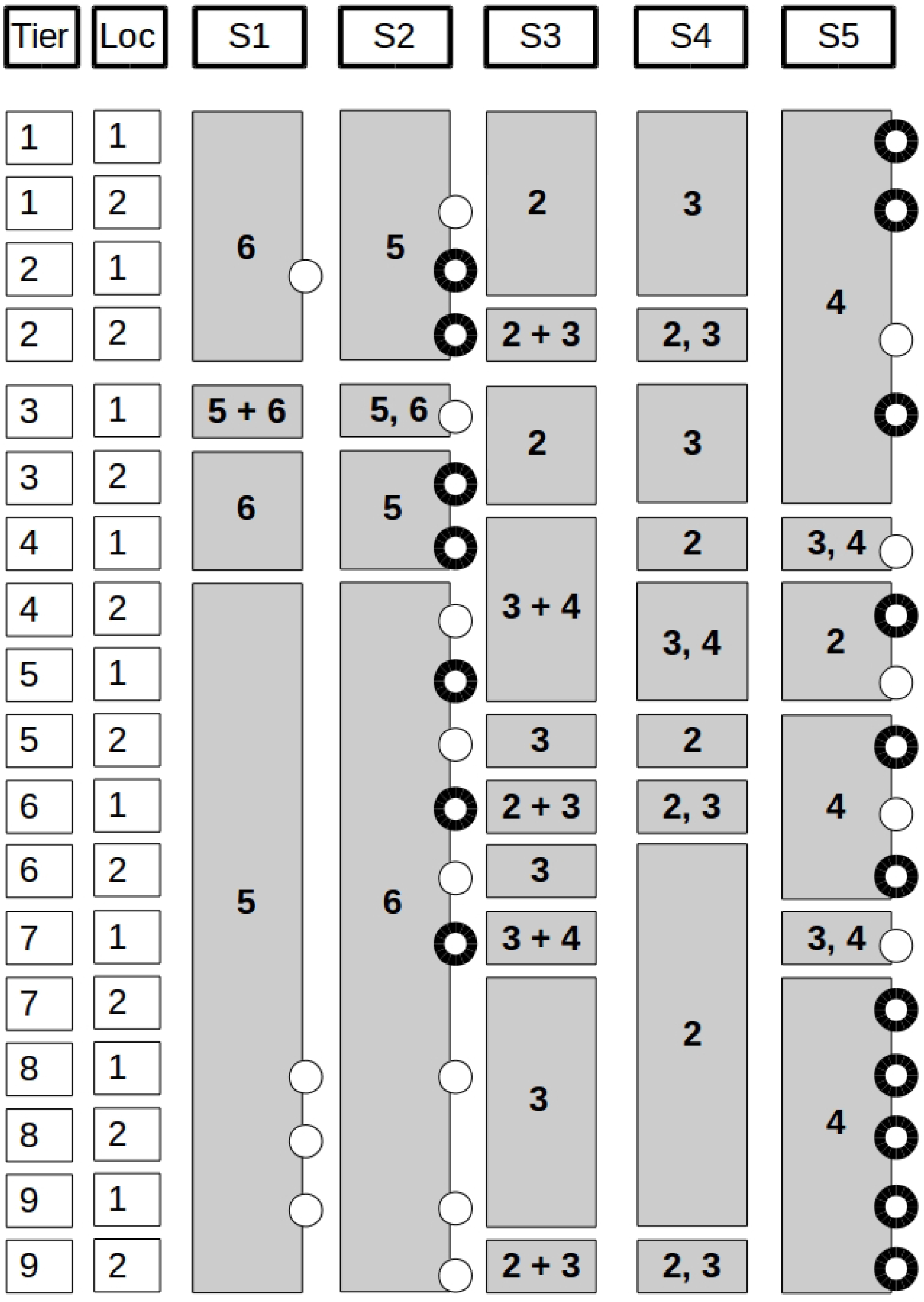
K-means clustering in the nine discriminant function tiers (1 to 9 from the uppermost to the lowest shore) in the two localities (1: Portocelo, 2: Corrubedo). S columns correspond to each of five cluster partition steps (the first resulting in two clusters and the fifth on six clusters; the last, sixth step is not shown as it results in the same result of seven single-variable clusters). The grey boxes show the measure(s) leaving a cluster in each step. The “*i+j*” code indicates that variables *i, j* were in the same new cluster, and “*i,j*”, that two new single-variable clusters were created, one including *i* and the other *j*. In all cases, S5 resulted in a cluster including measures 1 and 7. The circles mark partitions showing the maximum BoCluSt support (i. e., allocation variance = 0). Bold circles indicate the partition showing the maximum support in a thinned BoCluSt execution in that tier (see Materials and Methods for explanations).

The results for the variance of eigenvalues and average (non partial) correlation between variables were very similar for the two localities (Figure 4). Two remarkable results were obtained: First, the upper-shore phenotypes had clearly higher integration levels than the lower shore ones. This was a difference in the distribution of the variation in the seven measures, not (at least obviously) in overall variability: The average coefficients of variation were 0.153 and 0.320 in the most upper shore-like tiers and 0.207 and 0.165 in the most lower shore-like tiers in Corrubedo and Portocelo respectively. Second, there was a depression in integration levels for the intermediate section of the phenotypic gradient. The relationship between phenotypic tier and integration was clearly nonlinear: Introducing the quadratic term in the regression model resulted in marked increases in model fit and the quadratic regression coefficient was positive and significant. However, this coefficient reflects the fitted line's curvature and could be significant in the absence of a minimum in the middle of the phenotypic range. To confirm the existence of that minimum, we divided the range into three equal sectors (upper shore-like, intermediate, and lower shore-like phenotypes) of three tiers each, and made Welch two-sample, heterogeneous variances t tests comparing the sectors pairwise (as deviations from the locality average; three tiers and two localities per comparison).

**Figure 4.**
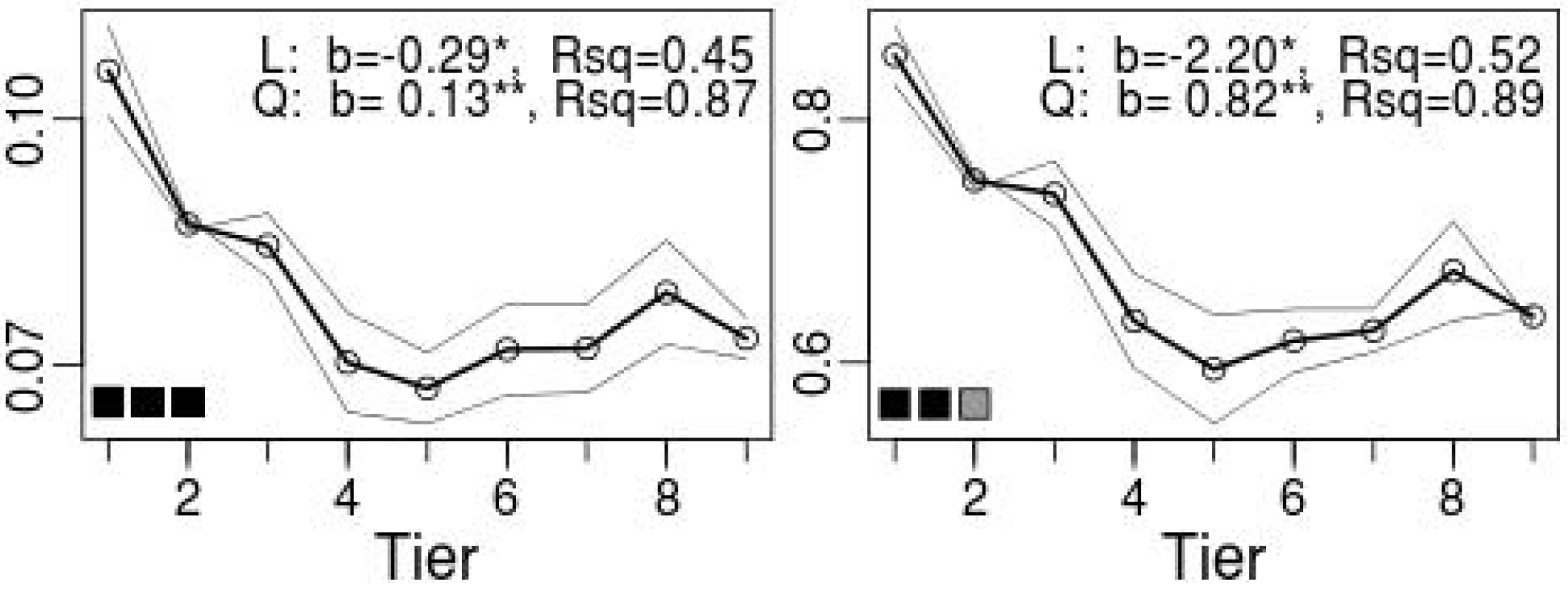
Overall integration in the nine discriminant function tiers (1 to 9 from the uppermost to the lowest shore). Left, variance of eigenvalues across the phenotypic gradient in the two localities (simple lines) and their average (bold line). Right, the corresponding results for average correlations between variables. L, results of linear regression analysis considering the tier number as covariable; b is the regression coefficient for this term. Q, results of a quadratic model considering the tier number and its square; b is the regression coefficient for the quadratic term. Rsq, r-square values for the two models. The square symbols correspond to the P-values of t tests comparing: left, the three first tiers with the three intermediate tiers; center, the three first and the three last tiers; right, the three intermediate tiers with the three last tiers. Black, grey and white squares correspond with P values < = 0.05, > 0.05 and <=0.10, and > 0.10 respectively.

The Newman and Girvan coefficient of modularity had clearly lower values for the upper shore-like shells, those with the highest overall integration (Figure 5). This could suggest that the reduced integrations observed in the intermediate and lower-shore-like tiers were due to increased modularity instead of an overall loss in data structure (see Figure 1). But Figure 5 shows that these reduced integrations were not associated with increases in within module correlation. Both the within and between module correlations were lower in the intermediate and low shore-like tiers, and it was only in a proportional sense that modularity increased, as correlations decreased more between than within modules. This correlation-based interpretation for the modularity results is supported by the close similarity between correlation quotients and modularity coefficients in Figure 5. Other measures of modular structure less dependent on correlations within and between modules, such as the Cheverud's index and the BoCluSt minimum variance criterion, failed to detect any clear trend across the phenotypic gradient.

**Figure 5.**
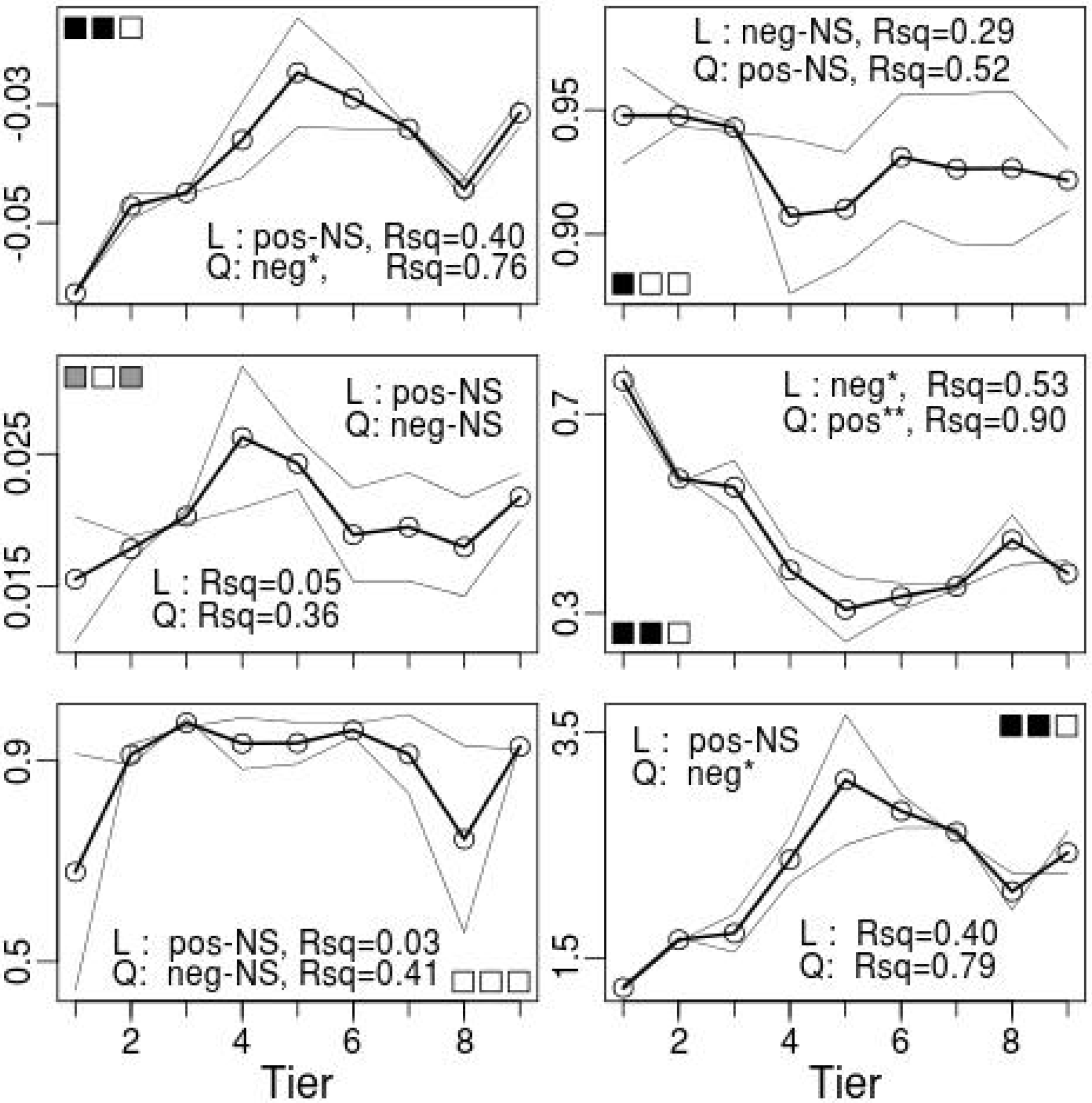
Modularity in the nine discriminant function tiers (1 to 9 from the uppermost to the lowest shore). Left column of graphs: Top, Newman and Girvan modularity across the phenotypic gradient in the two localities (simple lines) and their average (bold line); middle, Cheverud's index; bottom, one minus BoCluSt's minimum cluster allocation variance criterion. Right column: Top, average of the pairwise correlations between variables in the same cluster; middle, average of the pairwise correlations between variables in different clusters; bottom, quotient between the average correlations in the same and in different clusters. The regression and test results are represented as in Figure 3.

The partial correlations-based analyses reversed the trends found for those based on non-partial correlations (Figure 6). Now, the upper shore-like tiers showed the lowest levels of overall integration, as measured by the average partial correlation and weighted network connectivity, and, at the same time, the highest modularity. Again, this change in modularity was not due to a redistribution of the network link weights, as would be the case if they had increased within modules and decreased between modules. Both sets of link weights changed in the same direction, decreasing towards the upper shore-like tiers. As observed for the non-partial correlation results, the change in modularity was simply proportional, as the decreases in link weight tended to be proportionally larger within modules. The highest values for non-partial correlation modularities (Fig.5) and partial correlation overall integrations (Fig.6) tended to occur in consecutive non-extreme tier positions, which suggested real increases for these statistics in intermediate phenotypes. However, these high values not always resulted in significant differences, because their position did not coincide exactly with the three central tiers considered in the a priori designed t-tests.

**Figure 6.**
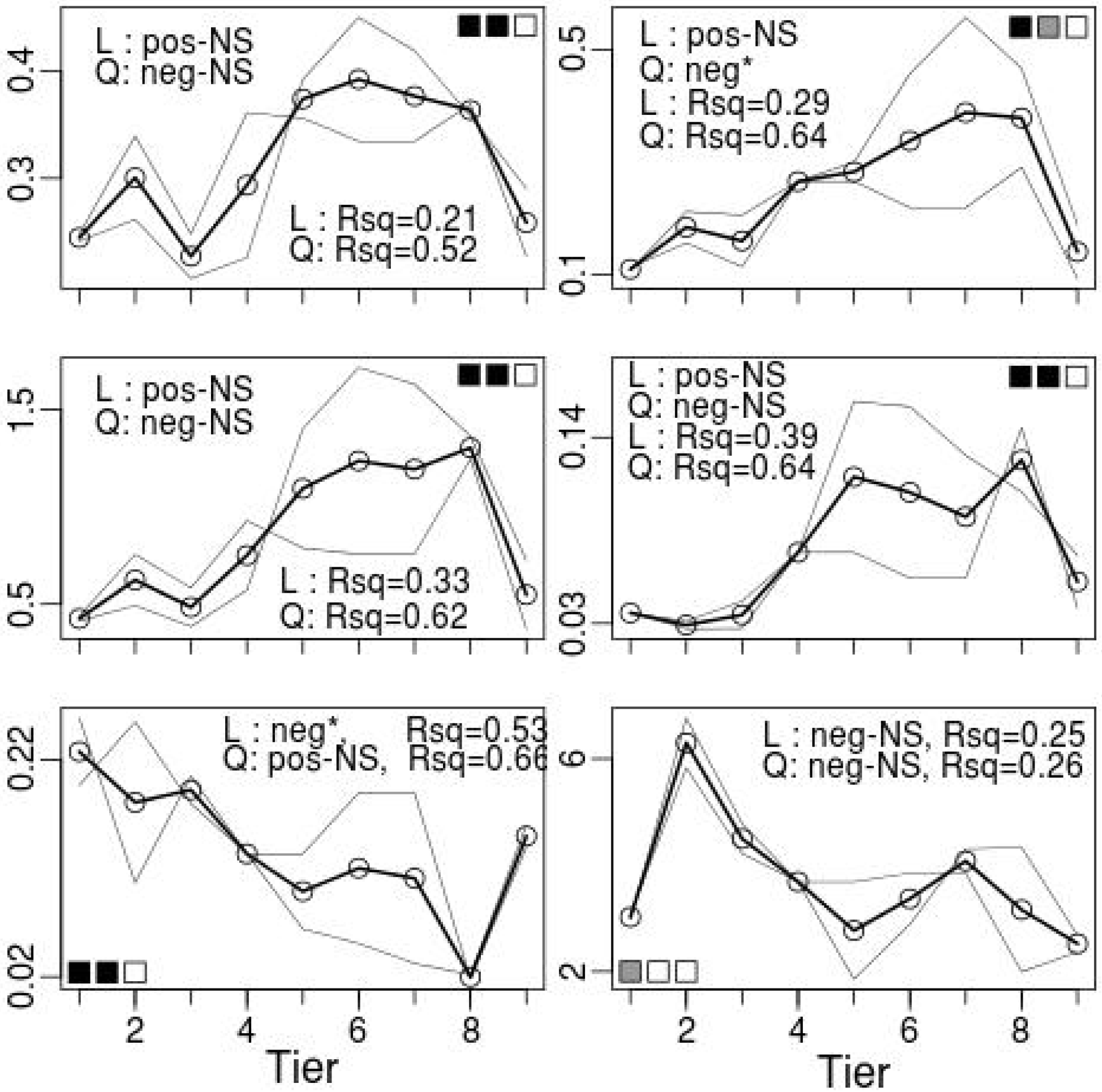
Overall integration and modularity based on partial correlations in the nine discriminant function tiers (1 to 9 from the uppermost to the lowest shore). Left column of graphs: Top, average absolute partial correlations between variables across the phenotypic gradient in the two localities (simple lines) and their average (bold line); middle, weighted network connectivity; bottom, Newman and Girvan modularity. Right column: Top, average of the absolute partial correlations between variables in the same cluster; middle, average of the absolute partial correlations between variables in different clusters; bottom, quotient between the average absolute partial correlations in the same and in different clusters.

The analysis of the measure-wise average non-partial correlations found markedly differentiated behaviors for measures 5 and 6 (Figure 7, top). Measure 5 made the major contribution to the progressive decrease in integration from the upper shore-like to the lower shore-like tiers, and measure 6 to the decreased integration seen for intermediate tiers. Figure 7 shows also that measure 5 had in average the lowest correlations in the lower shore-like shells and measure 6 the lowest in the upper shore-like ones, which was already reflected in the cluster analysis in Figure 3. In the lower shore-like tiers, it was measure 5 that left the initial group of measures in the first cluster partition, whereas measure 6 did so in the upper shore-like tiers.

**Figure 7.**
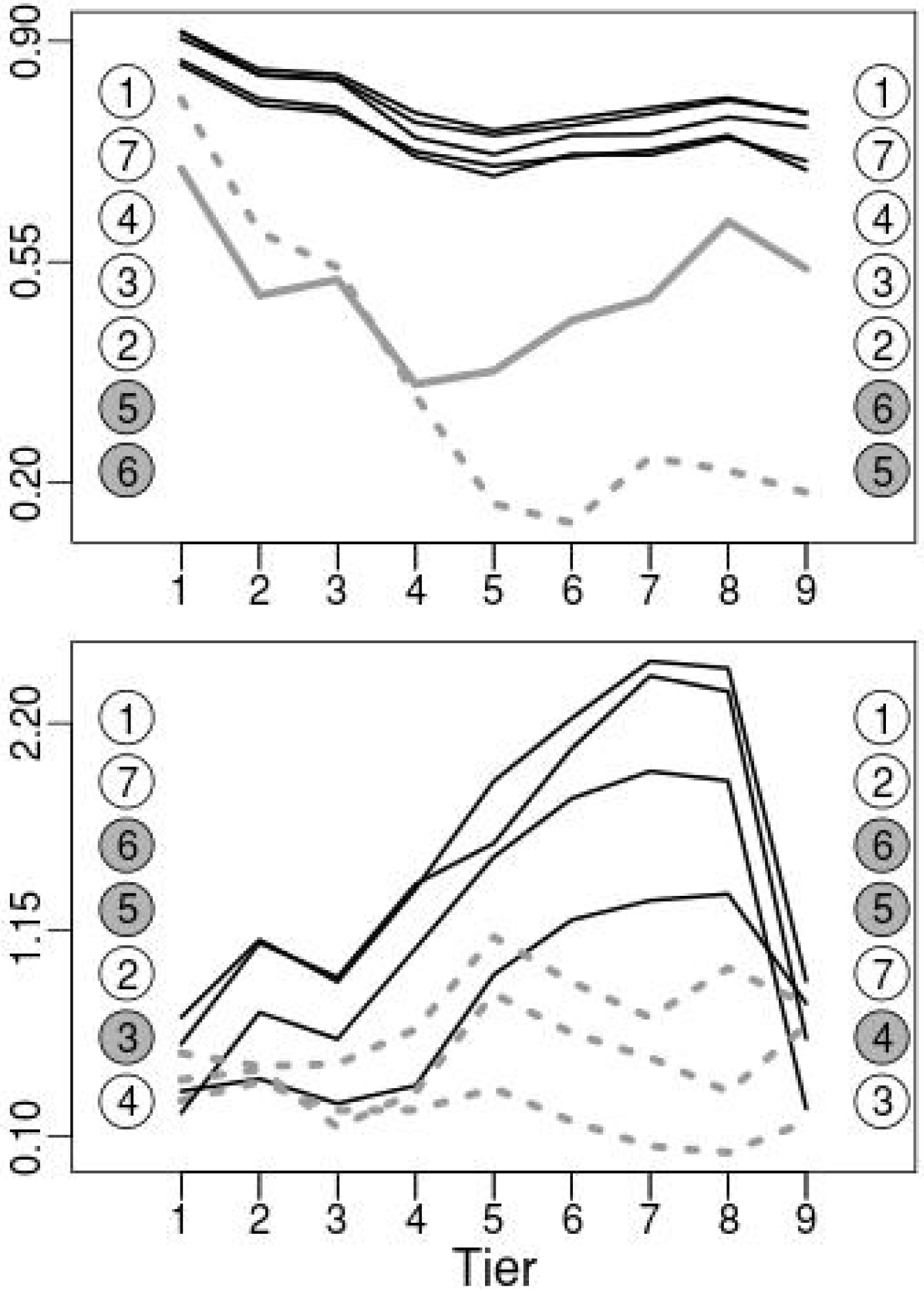
Average correlations (top) and edge strengths (bottom) for the seven shell measures in the nine tiers ( 1 to 9 from the uppermost to the lowest shore) for the discriminant function values. The measures' ranks for the first and last tiers are shown in the graphs' margins. Black lines are used for measures showing the main pattern of variation and dashed gray lines (continuous and dashed in the top graph to improve visibility) for measures diverging from this pattern.

**Figure 8.**
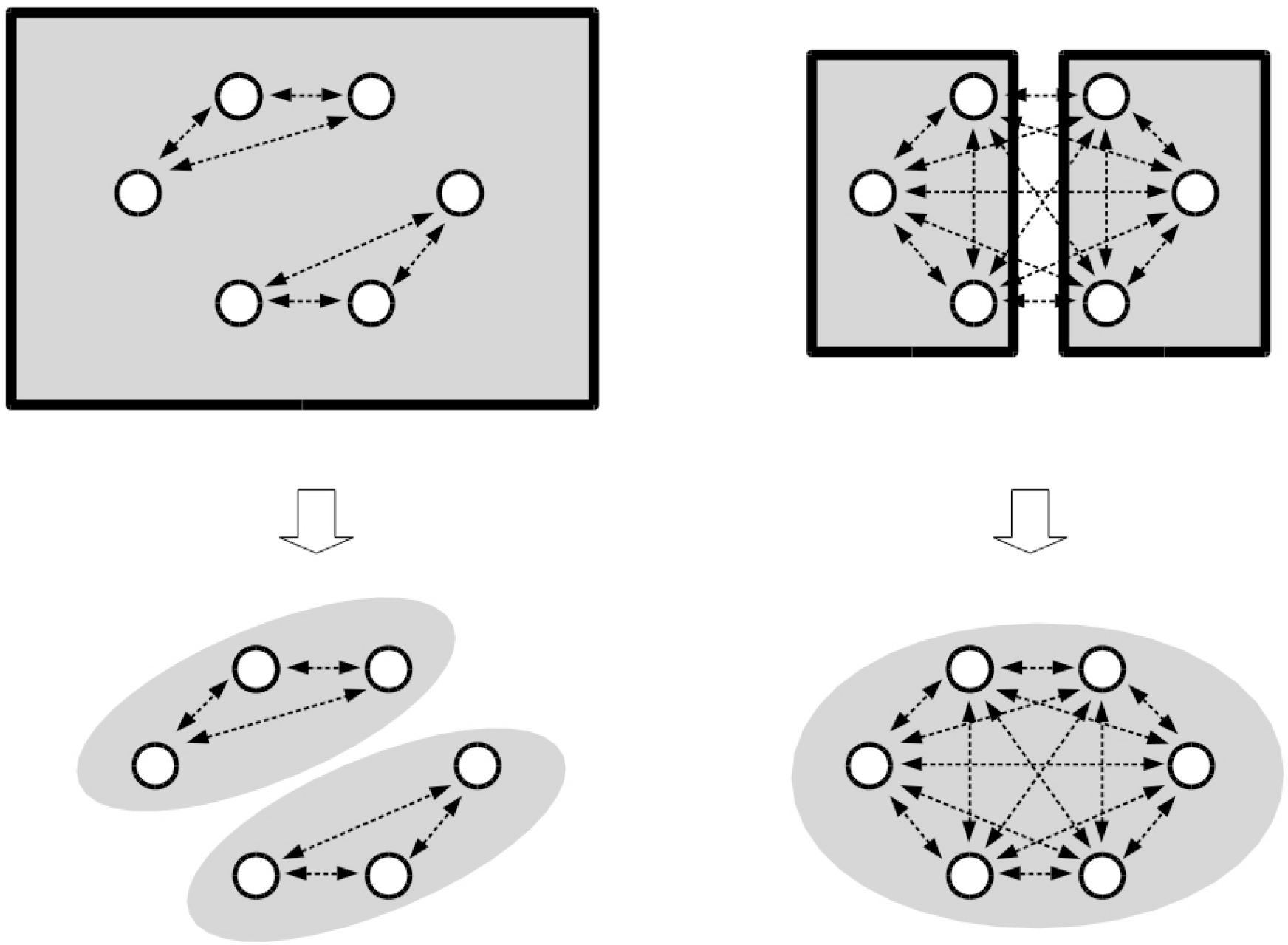
Schematic interpretation of results. Top, six measures (circles) are under the overall influence of one (left) or two (right) controlling variables. The ranges of these correlation-generating influences are indicated by rectangles. There are also weaker, pairwise sources of correlation (partial correlations; line arrows) independent of the controlling variables. An analysis using non-partial correlations as link weights would detect at right two communities (grey areas), less overall integration and higher modularity, whereas an analysis based on partial correlations (bottom) removing the effect of the controlling variables, would do so at left.

The logistic regression analysis of viability after transplant between shore levels found (see Appendix 1 for the full results) significant coefficients for shell measure 2 (b = 1.218, P = 0.017), measure 5 (b = 0.711, P = 0.017) and measure 6 (b = 0.585, P = 0.007), with no significant crossproducts or correlational selection coefficients. However, because of the considerable multicollinearity in this analysis (see the correlation matrix in Appendix 2), we made a complementary, stepwise logistic regression attempt to fit a more compact model to the data. The resulting model included the transplant destination x measure 5 crossproduct (b = 0.040, P = 4.3 e-6), corresponding to divergent selection, measure 2 (b = 0.026, P = 0.029,), and the transplant destination x measure 1 x measure 2 (b = 0.059, P = 0.042) triple crossproduct corresponding to divergent correlational selection. It must be noted that all positive crossproduct coefficients were consistent with adaptation, because shell measures had larger means in the upper shore-like snails due to these snails' larger overall size, and the transplant destinations were coded 5 to 1 from highest to lowest shore level. Thus, positive crossproducts corresponded to higher viability for upper shore-like shell measures or pairs of measures in upper shore destinations and for lower shore-like shells in lower shore destinations. Because all phenotypic correlations were positive, the positive significant coefficients observed for the two significant crossproducts in the stepwise regression would support an evolved association of phenotypic correlations with correlational selection. However, no relationship was found between phenotypic correlations and correlational selection coefficients in the complete model ( r = -0.020, P = 0.932, 19 d. f.) or between these correlations and divergent correlational divergent coefficients (r = 0.005, P = 0.982, d. f. = 19). Restricting the calculations to the regression coefficients having the lowest P-values did not narrow this relationship (not shown). Similarly, the regression coefficients for crossproducts involving measures 5 and 6, whose phenotypic correlations with the remaining measures were the lowest in average and also showed the steepest changes across the phenotypic gradient, were not clearly different from those of the other measures (Appendix 1).

However, the multicollinearity in the logistic regression model would again render the estimated coefficients unstable and difficult to interpret. We tried to get an additional view of the fitness impact of each shell measure by carrying out separate analyses of models including one shell measure, the transplant destination and all crossproducts involving that measure (measure x destination, measure x rest of measures, and triple cross products). The McFadden's pseudo-R squared (Frsq) values were, for measure one to seven, 0.0429, 0.0346, 0.0352, 0.0475, 0.0527, 0.0386 and 0.0409, measure 5 holding the first rank and measure 6 the fifth. While measure 5 held the first rank its value was not much higher than the rest. Thus, no clear support was found for a measure-wise relationship between phenotypic correlations and selection pressures in this hybrid zone.

However, some evidence for an overall relationship between selection and phenotypic integration was found in separate analyses of the extreme shore levels. Frsq was higher when the complete-model logistic regression was applied to a data set containing only the snails (of any shell type) transplanted to the highest shore level (level 5, Frsq = 0.237) than to a data set containing only snails transplanted to the lowest shore level (level 1, Frsq = 0.153). The same situation was found when level 5 was pooled with level 4 (Frsq = 0.125) and level 1 was pooled with level 2 (Frsq = 0.088), and also when the calculations were made separately for each location (Portocelo, Frsq = 0.356 and 0.199 for levels 5 and 1, and 0.151 and 0.131 for the pool of levels 5 and 4 and that of 1 and 2 respectively; Corrubedo, Frsq = 0.214 for the pool of levels 5 and 4, and 0.126 for that of levels 1 and 2; it was not possible to obtain a separate value for Corrubedo level 1 because this regression had a residual deviance = 0 and Likelihood = 1; this was also the analysis involving the least number of recaptured snails). These results were consistent with more restrictive selection on shell shape in the upper shore, where integration levels were highest.

## Discussion

The integration comparisons found remarkable differences between ecotypes, not detected in univariate analyses of the same traits in this hybrid zone. These differences would condition the future evolution of the ecotypes and the whole hybrid zone, as integration is expected to affect evolvability and to constrain the multivariate directions of selection response (Eroukmanoff and Svensson 2008). These differences between ecotypes did not correspond to differences in the amount of variation, either phenotypic (see above) or in terms of genetic markers (no marked differences in heterozygosity for AFLP markers or nuclear and mitochondrial sequences, as seen in Butlin et al. 2014 supplementary Table 2). Thus, there was no evidence of changes in developmental noise, canalization (Waddington 1942) or “genetic robustness” (Armbruster et al. 2009). Neither did the differences simply consist in uniform changes in the overall correlation between variables, but in changes in the structure of the correlation, because the composition of the identified variable modules changed across the gradient of shell phenotypes.

The upper shore ecotype showed higher integration when using non-partial correlations. The size of the difference was considerable in terms of average absolute correlation, as it went from about 0.8 to 0.6 from the upper shore to lower shore ecotype. According with common expectations about the evolution of integration, the higher integration in the upper shore would have been the consequence of stronger correlational selection, such as found in endemic or ecologically specialized versus generalist, or ancestral versus derived, species. This interpretation would be supported in this hybrid zone by the better fit obtained by natural selection regression models when applied to snails transplanted to the upper shore, and also by the shore distribution of the two ecotypes. While some upper-shore like snails are occasionally observed even at the lowest shore levels, the lowest shore-like ones are never found in the highest upper shore (personal observation). This points to an stronger selection on shell shape in the upper shore. There, the dryness, sun exposition and presence of predatory crabs could require special multi character adaptations by snails (see Mikolajewski et al 2015 for an example of increased morphological integration in predation-exposed dragonfly larvae). In fact, the gastropod community is impoverished in this area. Besides of limpets, only the snail *Melarhaphe neritoides* can be found at the upper shore limit of the *Littorina saxatilis* distribution (personal observation). The differences between ecotypes could also meet the expectation that derived, more specialized populations show higher integration than the ancestral ones, because the upper-shore ecotype could be in fact the derived one. Butlin et al. (2014) found evidence that the two ecotypes diverged in situ after colonization of the Galician shores at the end of the last glaciation, and speculate that *P. marmoratus*, the crab whose predation this ecotype would be adapted to resist, colonized these shores later, given that it is a warmer climate species. At present, this possibly derived condition is not associated with a marginal geographical distribution. Snails with the same shell phenotype can be found in large populations in the extensive sheltered bays and rias in Galicia, whereas the lower shore ecotype is restricted to very exposed areas. Such derived condition would not give clear-cut support to the expectation that functional specialization leads to the evolution of variationally independent modules (Wagner et al. 2007), at least within the studied character set. The upper shore-like phenotypes had the lowest Newman and Girvan modularity when considering non-partial correlations, and their higher modularity when considering edge strengths was accompanied by weak links within modules. Most of the difference in integration between ecotypes was generated by changes in correlations involving measure 5, but the functional role of this measure on resistance to predation or dry conditions is not obvious. Perhaps measure 5 is correlated with some unmeasured shell variable, as for example thickness, or ridge number or depth.

In case the upper shore ecotype had indeed differentiated in the sea-exposed areas, it could be interpreted that their upper shore adaptations enabled it to invade the more sheltered shores, also inhabited by *P. marmoratus*, and by additional predatory crab species such as *Carcinus maenas* (Linnaeus 1758). This would constitute an example of association between phenotypic integration and invasiveness. The relationship between these two features is still not too well understood, however.

Godoy et al. (2012) and Osunkoya et al. (2014) found that invasive plant species had, albeit to a limited extent, higher integration than related native species, and Colautti and Barrett (2011) found considerable integration in the invasive plant *Lithrum salicaria* L., as strong genetic correlations among life-history traits constrained the divergence among invading subpopulations of this species.

The cluster analysis detected consistent differences in clustering order, and therefore correlation structure between the extreme tiers in the phenotypic range (analogous differences between the two ecotypes had already been found in comparisons of matrix orientation and shape in Garcia 2012) and instability in this order in the intermediate phenotypic tiers. This instability corresponded to a significant depression in phenotypic integration as measured by the eigenvalues' variance, and a nearly significant one for the average correlation (Pavlicev et al. 2009 found that the absolute average correlation consistently underestimates integration). Given that a large proportion of phenotypic variation for shell measures has a genetic basis in this species (Conde-Padin et al. 2009), the depression in phenotypic integration observed for the intermediate phenotypes supports the view that hybridization could result in the release of new multicharacter genetic variation and thus open new evolutionary trajectories. This was not related with transgressive segregation for individual traits, but with changes in the correlation matrix. It is interesting to note that, in contrast with the expected consequences of genetic incompatibilities between parents (Fitzpatrick 2008) the disruption of regulating or developmental processes possibly causing these changes in integration was not associated with marked fitness reductions for snails with intermediate phenotype. However, Cruz and Garcia (2001) found some reductions in embryo number and size in intermediate females, and Sa Pinto et al. (2013), a higher degree of DNA fragmentation in the sperm of intermediate males, associated with increased embryo abortion rates and reduced adult viability.

The observation that measure 6 had a dominant role in the integration depression and that correlations involving measures 5 and 6 showed a clearly differentiated evolution across the hybrid zone suggest different sources for the correlations in this study. While the results involving measures 1, 2, 3, 4 and 7 were rather consistent with the stability expected for pleiotropy-generated correlations (Klingenberg 2008), those involving measures 5 and 6 were more labile and likely related with linkage disequilibrium. However, this interpretation is simply speculative, given that phenotypic correlations might not reflect the underlying genetic architecture (von Dassow and Munro 1999, Armbruster 2014).

The hypothesized role of past correlational selection in shaping the present correlations was not supported by the relationship between correlational selection coefficients and phenotypic correlations in these data. However, the failure to find a relationship between phenotypic correlations and multiple regression-estimated selection coefficients could be the consequence of the high multicollinearity in the regression models applied, which would have prevented to estimate regression coefficients in a reliable and stable way. A different-approach analysis, based on principal components, (Garcia 2014b) found four principal components (PCs) significantly affected by divergent selection. The PC most affected was the second, which consisted essentially in a contrast of measure 5 with measure 6 (and to a less extent, also with measures 1 to 4 and 7) followed by the first one, which had positive coefficients for all measures and thus likely represented general shell size. The observation that the second PC, contributing with 7.15% of total variance, was more affected by divergent selection than the first one, accounting for 85.31%, underlaid the conclusion that the correlation matrix was not aligned with the vector of divergent selection. Thus, these results do not fully fit to the view that integration could constitute an adaptation increasing the correspondence between the individuals' phenotypic and environmental patterns of variation (Armbruster et al. 2014). However, this did not preclude any relationship between selection and correlation. The finding that it was the PC contrasting measures 5 with the rest, mostly measure 6, that suffered the most divergent selection was consistent with the large change across the phenotypic range (i. e., more positive combinations for upper shore-like shells and less positive for lower shore ones) for correlations involving measure 5. In addition, the first PC, the next most affected by divergent selection, had positive coefficients for all shell measures, which implied that large and small shells were more viable in the upper and lower shore respectively. This could have reinforced the presently observed strong positive correlations, specially between measures 1 to 4 and 7, no seriously affected by selection acting on the second PC. However, it must be taken into account that estimates of selection pressures used here were obtained in a short term experiment in which adult animals were transplanted to unfamiliar environments. It was not designed to provide a detailed description of natural selection pressures in the long term in each shore level. The present study supports the view that, despite the considerable stability in integration patterns commonly observed across species (Young 2004, Marroig & Cheverud 2005, Goswami 2006, Sears et al. 2013), these patterns can diverge in evolutionary spans short enough to generate differences within the genus level (Kolbe et al. 2011, Sanger et al. 2012) or even within species, as in this hybrid zone, under the appropriate selection pressures.

The contrasting results found for the variation in overall integration and modularity when using non-partial and partial phenotypic correlations points to different, overlapping levels of regulation of shell shape. Some controlling variables acting on several shell measures simultaneously could contribute to the measured non-partial phenotypic correlations. The changes observed in the corresponding partial correlations would be due to the removal of these controlling variable effects. The remaining, independent sources of correlation could operate in the reverse direction (Figure 7), thus generating the increased average partial correlations for several not-extreme phenotypic tiers. Indeed, hybridization could cause both increases or decreases in integration. For example, increases could result if some regulating genes from different origins remained linked after hybridization (Wagner et al. 2007), and decreases if they did not, or as a consequence of disruption of some regulation circuits such as those controlling gene expression by different enhancers or transcription factors (Wagner et al. 2007). The constrasting patterns of variation for the two kinds of correlations, integration and modularity measures across the phenotypic gradient point to differences in the mechanisms regulating shell development in the two ecotypes in this hybrid zone.

This study has found large differences in integration between the two ecotypes in this hybrid zone. Given that these ecotypes could have differentiated in situ, according to a parapatric model (Quesada et al. 2007, Butlin et al. 2014), our results would show that changes in integration can occur in a short evolutionary time and be maintained in the presence of gene flow, and also that this gene flow may reduce phenotypic correlations among characters in hybrids, potentially lessening genetic constraints on evolutionary trajectories.

